# Avian recolonization of unrestored and restored bogs in Eastern Canada

**DOI:** 10.1101/2021.11.26.470119

**Authors:** André Desrochers, Line Rochefort

**Affiliations:** Département des sciences du bois et de la forêt, Université Laval, Québec, Canada; Peatland Ecology Research Group and Centre for Northern Studies, Université Laval, Québec, Canada

**Keywords:** Peatland, Ecological Restoration, Boreal, Ornithology, Occupancy, Species Diversity, Habitat Selection, Autonomous Recording Units

## Abstract

Over the last several decades, peat has been extracted from bogs of temperate, populated regions of Eastern Canada, leaving large areas devoid of vegetation if unrestored. For the last 25 years, projects have been conducted in these regions to re-establish vegetation and facilitate recolonization by wildlife. We tested whether vegetation structure and bird species assemblages 10 to 20 years post extraction differ among natural, unrestored and restored bogs at the scales of individual sites and entire bogs. We conducted bird counts and vegetation surveys between 1993 and 2019, using both point counts (309 sites) and Autonomous Recording Units (80 sites). According to our vegetation surveys, restoration of sites that were previously used for peat harvesting accelerated the establishment of *Sphagnum* and herbaceous strata, but ericaceous and tree strata were unaffected over a 17-year period. None of the bird species with large home ranges were associated specifically to natural, unrestored, or restored areas at the bog level. Bird species diversity was similar in restored and natural sites, but lower in unrestored sites. Alder Flycatcher and American Goldfinch occupied restored and unrestored sites more frequently than natural sites, independent of the number of years post extraction. Occupancy of restored sites by Palm and Yellow-rumped Warblers increased over the years, reaching levels similar to those in natural sites 20 years after restoration was implemented. Occupancy of restored sites by Song and Savannah sparrows increased from 1993-2019 and diverged from their declining occupancy of natural sites. Species assemblages of restored and unrestored sites differed significantly from those of natural sites soon after peat extraction ceased or post restoration. But assemblages from restored and unrestored sites became progressively similar to those of natural sites during the first 20 years, especially in restored sites. We conclude that bird species assemblages of restored bog sites are converging toward those of natural sites, and that restoration provides novel habitats for regionally declining species, e.g., Savannah Sparrows.

## Introduction

Sphagnum-dominated peatlands, or bogs, sometimes dominate boreal and subarctic landscapes (Gorham, 1990) and extend south to temperate latitudes, with increasing isolation and distinctiveness from landscapes (Brewer, 1967; Calmé and Desrochers, 2000; Poulin and Pellerin, 2001). In temperate, populated regions, they are often drained for the extraction of peat and other uses. In Eastern Canada, horticultural peat extraction has occurred for nearly a century, starting with small-scale manual extraction by blocks, which was gradually replaced by large-scale extraction with vacuum-harvesters pulled by tractors (Girard et al., 2002). A small proportion of peatlands has been used for peat extraction in boreal Canada (<0.03 %; CSPMA (2014)), but undisturbed peatlands, especially large ones, have become rare in populated areas of Eastern Canada (Pellerin, 2003; Poulin et al., 2016). Since the 1990s, wetland ecologists have been aware of the potential impact of peat extraction on regional biodiversity, and the potential of ecological restoration (Wheeler and Shaw, 1995; Gorham and Rochefort, 2003). During the same period, the peat moss industry collectively decided to develop ecological restoration methods to facilitate the return to peat-accumulating ecosystems, with associated flora and fauna (Rochefort, 2000).

Wetland restoration is challenging, especially in colder climates where ecosystem processes are slower (Moreno-Mateos et al., 2012). Early on, some researchers thought that bogs formerly subjected to peat extraction may revert to peat-accumulating ecosystems without active restoration (Green, 1983; Smart et al., 1989; Famous et al., 1995). After decades of bog restoration and comparisons with unrestored peat-extracted bogs, this hypothesis is no longer supported (Poulin et al., 2005; Lavoie et al., 2003) and consequently ecological restoration of peat-extracted bogs is taking place in several northern countries (Andersen et al., 2017; Chimner et al., 2017; Karofeld et al., 2016). But we do not know yet whether restoration can produce wildlife species assemblages similar to those of natural sites.

Measuring the success of ecological restoration is not a trivial matter, and should fulfill several criteria, some of which include reaching a desired species diversity, assemblage and indigenous species (Prach et al., 2019). The return of plant species assemblages and vegetation structure has been the focus of earlier ecological restoration work (Young, 2000; Hugron et al., 2020), some of which was motivated by the ‘if you build it, they will come’ assumption (Palmer et al., 1997). Restoring vegetation may be necessary, but insufficient however, for a lasting recolonization of restored sites by wildlife, as exemplified by several studies (e.g., Cross et al., 2019). In the case of bogs, previous work by Desrochers et al. (1998) showed substantial differences among bird species assemblages in unrestored vs. natural sites, but their work was based only on surveys spanning 3 years, at a time when bog restoration was in its infancy, with few recovery years since restoration.

This study documents the re-establishment of vegetation and associated breeding birds in Québec and New Brunswick, from 1993 to 2019. More specifically, we compared vegetation cover and site occupancy by songbirds between natural, unrestored, and restored sites. Additionally, we tested whether the occupancy of species with large home ranges was associated to aggregated areas of undisturbed, unrestored and restored sites at the scale of entire bogs.

## Study Area and Methods

We sampled bird populations in 38 open raised bogs of Québec and New Brunswick, sometimes hundreds of kilometers apart (Fig. 1). Each year of data collection, we sampled natural, unrestored (without management post extraction), and restored sites in most of the bogs. Throughout this paper, we define sites as specific, contiguous areas of bogs with a homogeneous class (natural, restored, etc.) We assume that our sampling approach led to no major confounding effects of regional and temporal variation among bogs.

**Figure 1:**
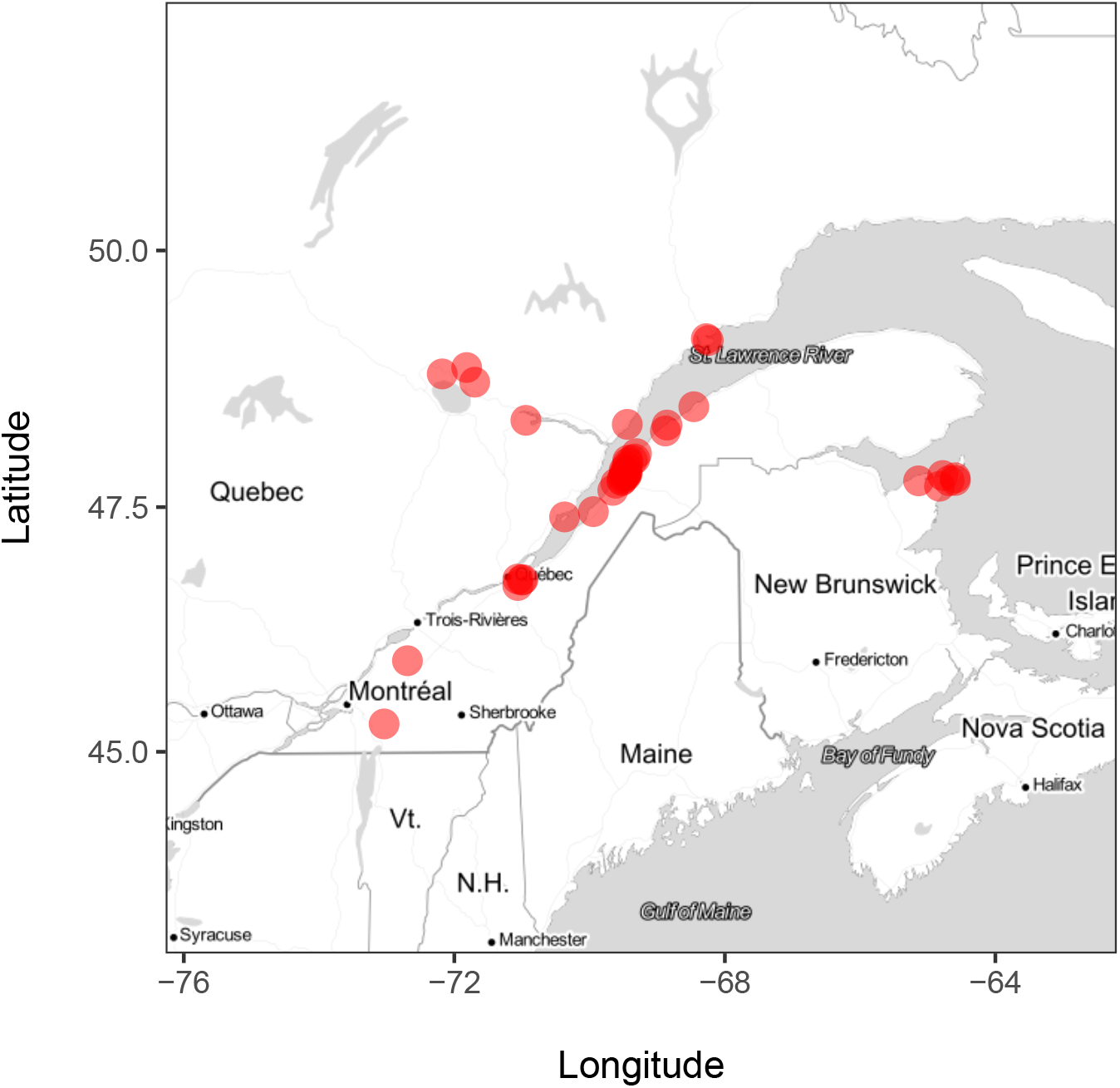
Study area.

### Bird Point Counts

We performed 1659 point counts at 309 sites in 1993, 1996, 1999, 2002, 2005, and 2011, in the absence of strong wind or rain (Table 1). Point counts were separated by > 200 m and performed between 20 May and 15 July. Each point count site was sampled on 1 to 5 occasions within a given breeding season. Point count duration was 10 min in the mornings and 5 min in the evenings, with the exception of 15-min point counts with playbacks of alarm and mobbing calls (details in Corbani et al. (2014)), which were performed in 2011 for a separate study. All 12 observers who conducted point counts had > 5 y of birding experience and were assigned natural, unrestored and restored sites. We assume that variation among observers did not create bias.

**Table 1:** Numbers of bird surveys by method and time of day.

| Protocol | Time | Minutes | Year |  |  |  |  |  |  |  |
| --- | --- | --- | --- | --- | --- | --- | --- | --- | --- | --- |
|  |  |  | 1993 | 1996 | 1999 | 2002 | 2005 | 2011 | 2018 | 2019 |
| Passive Point Count | Evening | 5 | 39 |  |  |  |  |  |  |  |
|  |  | 10 | 23 |  |  |  |  |  |  |  |
|  | Morning | 5 | 17 |  |  |  |  |  |  |  |
|  |  | 10 | 376 | 96 | 210 | 203 | 325 | 142 |  |  |
| Point Count with Playback |  | 10 |  |  |  |  |  | 112 |  |  |
|  |  | 15 |  |  |  |  |  | 116 |  |  |
| Autonomous Recording Unit | Evening | 5 |  |  |  |  |  |  | 51 | 45 |
|  | Morning | 5 |  |  |  |  |  |  | 188 | 175 |

### Recordings

In 2018 and 2019, we deployed Wildlife Acoustics Song Meters (models SM3 and SM4; Wildlife Acoustics Inc., Concord, MA, USA) on 80 sites separated by > 200 m (Table 1). We programmed each of those Autonomous Recording Units (ARUs) to record one 5-minute sample per hour during deployments that lasted between 0.4 and 26 d (mean = 7.8 d). After deleting samples with excessive noise due to wind, rain, or malfunction, for each site, each year, we sampled two to five 5-min samples between 4:00 – 9:00 EDT (mean = 4.5 samples) on different days, and one to four samples between 19:00 – 22:00 EDT (mean = 1.2 samples), also on different days. Each recording was encoded on two channels with a 24 KHz sampling rate and a 16 bit resolution. Each recording sample was listened to with headphones by A.D. and all species heard were transcribed into an eBird list (Sullivan et al., 2009).

### Bog land cover

We measured the temporal evolution of land cover in bogs from a combination of aerial and satellite images taken throughout the study period, as well as GPS surveys. All spatial data were recorded as adjacent polygons and incorporated in an ArcGIS geodatabase (ESRI, 2019) using a decimal degree, WGS84 (EPSG 4326) projection. We ascribed one of the following categories to each polygon: natural, peat-extracted, unrestored, restored, pond. Sites not corresponding to those categories were ignored. We estimated the year of the ending of peat extraction and restoration for each polygon for which one of those categories occurred at least once. We updated bog land cover categories for each survey year. Sampling plots for vegetation and birds (see below) were characterized as unrestored (peat extracted), restored or natural only if > 50 % of their area corresponded to one of those categories. We retained only sites unrestored or restored 10 to 20 y before the surveys, because older unrestored sites had no restored counterparts with similar ages. We assumed that restored and unrestored sites less than 10 y old were not sufficiently different to greatly influence bird occupancy.

### Vegetation surveys

Prior to each point count, observers estimated the percent cover of bare peat, *Sphagnum*, herbaceous plants, ericaceous plants, shrubs (< 5 m) and trees, all on circular plots delimited with surveying flags and < 100 m from the observer. Strata sometimes overlapped, leading to total cover exceeding 100 %. Standard deviations among estimates on the same year for single sites ranged from 5.7 – 17.8 %. We did not perform vegetation surveys in 2018 or 2019.

### Statistical analysis

Traditional analyses of species occurrences were performed without accounting for imperfect species detection, e.g. in our former study of peatland birds (Desrochers et al., 1998). This basic approach not only precludes actual estimate of true site occupancy, but it may lead to biases if the detection process is dependent on factors associated with sampling time or location (Mazerolle et al., 2005). Repeated bird surveys at the same sites during a single breeding season, as in the present study, allow the estimation of detection probabilities, given that species are often encountered at certain surveys but not others. Assuming that species were invariably present or absent at given sites during the entire season each year (the closure assumption), it is possible to estimate detection probabilities and adjust estimates of site occupation accordingly (MacKenzie et al., 2002). Mackenzie et al’s approach use maximum-likelihood estimation to simultaneously model detection probabilities against covariates that may affect detection (time of day, etc.), and occupancy itself, against site characteristics.

We measured site occupancy by birds by accounting for imperfect detection with Mackenzie et al’s (2002) occupancy modelling. We conducted analyses separately for species with small home ranges (most songbirds) and species with large home ranges or usually detectable from distances exceeding 100 m (raptors, swallows, shorebirds, etc.). We considered species as significantly responding to natural, unrestored or restored sites when the weight of evidence of the null model was less than 5 % divided by the number of species tested, to maintain Type I error rate (false positives) at *α* = 5 % despite multiple testing.

### Birds with small home ranges

We compared bird species observed from natural, unrestored and restored sites, with two approaches. First, we examined species diversity by computing species accumulation curves (Soberón and Llorente, 1993). Given the multiple ways to obtain cumulative species numbers (depending on the ordering of sites), we used a bootstrap procedure (Efron, 1982), i.e we sampled *n* sites with replacement, each 100 times, and computed the mean and standard error of the number of species for each *n*. We used a maximum *n* of 22 sites, because that was the number of sites of the least sampled site category (restored). Species diversities from 22 sites were compared with a randomization test, i.e. by comparing the observed differences to a random distribution of 10,000 simulated differences from shuffled site categories and expressing a p-value as the quantile of the observed differences relative to the random distribution.

Second, we conducted a series of single-species analyzes. We built a site-specific covariate matrix consisting of one record per site, with columns representing year of survey, and site status. We built four survey-specific matrices, each with one record per site and one column for each sample: Julian dates, times of day (evening, morning), count durations, and protocols (ARU, passive point count, point count with playback of mobbing calls). Then, for each species encountered at least 50 times in sites meeting the site selection criteria (see above), we built a matrix with one record per site per year, with one column for each sample, and cells containing presence/absence data.

We compared three occupancy models for species with small home ranges. To limit the number of competing models,each of them included the same detection covariates : Protocol + Duration + Time + Julian.

1. ~ Protocol + Duration + Time + Julian ~ Year {null model}
2. ~ Protocol + Duration + Time + Julian ~ Year + Status
3. ~ Protocol + Duration + Time + Julian ~ Year * Status

The first model incorporated only detection covariates, with a hypothesized temporal trend on site occupancy. We designed the second model to evaluate the additive effects of trend and status, i.e. fixed occupancy differences between natural, unrestored and restored sites. The third model evaluated a temporal change in the differences between occupancies of natural, unrestored and restored sites.

We ran the suite of models first with natural, unrestored, and or restored sites to get general patterns, and re-ran the models with unrestored and restored sites only, to test whether site restoration was associated to changes in species occupancy. To estimate species responses to site restoration (vs. unrestored) we used multimodel inference, with occupancy estimates for site status averaged between models, weighted by model AICc weight (Burnham and Anderson, 2002).

### Bird with large home ranges

We followed the same modelling approach as with species with small home ranges, with the following differences. The site-specific covariate matrix consisted of one record per bog, with columns representing year of survey, and log areas of peat-extracted, natural, unrestored, and restored sites. Logs were used because we believe that a fixed increment in area, say 1 ha, has greater importance in small areas than in larger areas. We built four survey-specific matrices, each with one record per site and one column for each sample: Julian dates, times of day (evening, morning), count durations, and protocols (ARU, passive point count, point count with playback of mobbing calls).

We compared seven occupancy models for species with large home ranges. ‘.’ refer to the same covariates as in the previous set of models.

1. ~ . ~ Year {null model}
2. ~ . ~ Year + Natural
3. ~ . ~ Year + peat-extracted
4. ~ . ~ Year + Unrestored
5. ~ . ~ Year + Restored
6. ~ . ~ Year + Ponds
7. ~ . ~ Year + peat-extracted + Natural + Unrestored + Restored + Ponds

Each model used the same detection covariates as in the models for species with small home ranges. Models were designed to evaluate a general trend in occupancy from 1993 to 2019, irrespective of bog characteristics (null model), and the isolated or additive effects of areas of each bog characteristic.

### Bird species assemblages

We compared bird species assemblages of natural, unrestored and restored sites, based on 41 species observed at least 10 times in sites selected for the single-species analyses. We used a multivariate approach based on `site x species` matrices: a dendrogram for visualisation, and a Mantel test for hypothesis testing (Legendre et al., 2015, and references therein). In both cases we used binary distance estimates for each pairwise site comparison, i.e. the proportion of species which were present at only one of the two sites. With the Mantel test, we tested whether the site x site distance matrix for species was correlated to the distance matrix for site status (natural, unrestored, restored). Two analyses were performed, one including the three status types, the other comparing only unrestored and restored sites. A Mantel correlation of zero is obtained when species assemblages are independent of site status. We performed 10 000 permutations for each partial Mantel test.

All data preparation and statistical analyses were performed with R and associated `tidyverse` and `aiccmodavg` packages (R Core Team, 2020; Wickham et al., 2019; Mazerolle, 2020).

## Results

### Bog land cover and vegetation strata

Land cover statistics include only bogs sampled in the current study, and may not be representative of the entire set of peat-extracted bogs in Quebec, New Brunswick or Canada. Within the sampled bogs, the cover area changed greatly between 1993 and 2019. In the sampled bogs, restored sites went from zero to 818 ha, but we observed a net decrease of natural sites and ponds of 15 % and a 81 % net increase of unrestored sites (Table 2).

**Table 2:** Net temporal change in the land cover of bogs within 2 km of sampling sites, 1993-2019. RDL stands for the Rivière-du-Loup peatland complex. Values are in hectares (as of 1993), with changes from 1993 to 2019 in parentheses.

| Name | Location | Natural | Ponds | Extracted | Unrestored | Restored |
| --- | --- | --- | --- | --- | --- | --- |
| Bagotville | 48.36 N, 70.94 W | 496.1 (0) | 0 (0) | 0 (0) | 109.6 (0) | 0.0 (0) |
| Bois-des-Bel | 47.97 N, 69.43 W | 185.5 (0) | 0 (0.1) | 0 (0) | 11.1 (-7.5) | 0.0 (7.4) |
| Cacouna-Station | 47.88 N, 69.45 W | 71.6 (0) | 0 (0) | 0 (0) | 86.8 (0) | 0.0 (0) |
| Coteau-du-Tuf | 47.97 N, 69.37 W | 2.5 (0) | 0 (0) | 0 (4.2) | 24.8 (-4.2) | 0.0 (0) |
| Grande Plée Bleue | 46.77 N, 71.07 W | 590.2 (0) | 15 (0) | 0 (0) | 0 (0) | 0.0 (0) |
| Ile-aux-Coudres | 47.41 N, 70.36 W | 71.5 (-20) | 0 (0) | 74.8 (-74.8) | 23.5 (94.8) | 0.0 (0) |
| Inkerman | 47.71 N, 64.81 W | 736 (-149.8) | 11.7 (-5.3) | 197.7 (136.9) | 65.2 (6.5) | 0.0 (11.6) |
| Isle-Verte | 48.03 N, 69.31 W | 27.3 (0) | 0 (0) | 0 (0) | 89 (0) | 0.0 (0) |
| Isle-Verte SW | 47.98 N, 69.33 W | 0 (0) | 0 (0) | 35.5 (-35.5) | 0 (35.5) | 0.0 (0) |
| L'Ascension Ouest | 48.73 N, 71.7 W | 1948.2 (-98.6) | 11.2 (0) | 109.6 (79.8) | 4.7 (-0.2) | 0.0 (19) |
| Lac Malobès | 48.25 N, 68.88 W | 0 (0) | 0 (0) | 0 (0) | 9 (0) | 0.0 (0) |
| Le Parc | 47.87 N, 69.49 W | 9.4 (0) | 0 (0) | 0 (0) | 36.1 (0) | 0.0 (0) |
| Les Escoumins | 48.32 N, 69.45 W | 605.8 (-66.3) | 7.6 (-2.7) | 105.8 (77.1) | 26.1 (-21.8) | 0.0 (13.8) |
| ND-du-Portage | 47.74 N, 69.61 W | 119.2 (0) | 0 (0) | 101 (-101) | 28.6 (101) | 0.0 (0) |
| Pointe-au-Père | 48.49 N, 68.47 W | 48.5 (0) | 0 (0) | 171.6 (-171.6) | 12.6 (148.9) | 0.0 (22.8) |
| Pointe-Lebel | 49.14 N, 68.26 W | 1762.3 (-358.2) | 198.2 (-5.6) | 610.4 (104.3) | 1.7 (205.9) | 0.0 (53.6) |
| Pokesudie | 47.81 N, 64.77 W | 113 (0) | 3.1 (0) | 81.1 (-81.1) | 58.3 (-31.3) | 0.0 (112.4) |
| RDL Centre | 47.8 N, 69.49 W | 7.4 (0) | 0 (0) | 0 (12.5) | 135.1 (-23.8) | 0.0 (0) |
| RDL Chemin-du-Lac | 47.76 N, 69.53 W | 26.5 (-12.5) | 0 (0.9) | 208 (-197.3) | 13.5 (151) | 0.0 (56.2) |
| RDL Chemin-du-Lac 2 | 47.78 N, 69.52 W | 0 (0) | 0 (0) | 70.4 (-13) | 0 (13) | 0.0 (0) |
| RDL Côte-Sud | 47.84 N, 69.47 W | 145.1 (0) | 0 (0) | 106.1 (-52.9) | 18.6 (51.6) | 0.0 (1.3) |
| RDL Président-Ouest | 47.79 N, 69.5 W | 363.6 (-131) | 0 (0) | 241.4 (12.5) | 47.7 (96.4) | 0.0 (22.1) |
| RDL St-Laurent | 47.82 N, 69.47 W | 271.3 (-70.9) | 0 (0) | 214.3 (134.3) | 371.9 (-82) | 0.0 (18.6) |
| RDL St-Laurent 2 | 47.82 N, 69.46 W | 3.9 (0) | 0 (0) | 59.7 (31.1) | 31.2 (-31.2) | 0.0 (0) |
| RDL St-Modeste | 47.85 N, 69.45 W | 0 (0) | 0 (0) | 52.3 (-5.3) | 26.8 (5.3) | 0.0 (0) |
| RDL St-Modeste 2 | 47.83 N, 69.45 W | 0.9 (-0.9) | 0 (0) | 45 (-44.1) | 0 (45) | 0.0 (0) |
| RDL St-Modeste 3 | 47.84 N, 69.48 W | 0 (0) | 0 (0) | 15.3 (-15.3) | 0 (15.3) | 0.0 (0) |
| RDL Verbois | 47.84 N, 69.45 W | 32.7 (-32.7) | 0 (0) | 170.4 (-35.5) | 8.2 (39) | 0.0 (29.3) |
| Rivière-Ouelle | 47.47 N, 69.93 W | 984.4 (-242.7) | 0 (0) | 480.5 (238.4) | 30.2 (-3.8) | 0.0 (4.4) |
| St-Alexandre | 47.66 N, 69.67 W | 49 (0) | 0 (0) | 266.3 (-266.3) | 68.8 (266.3) | 0.0 (0) |
| St-Arsène | 47.94 N, 69.43 W | 126.8 (0) | 0 (0) | 153 (-1.3) | 29.6 (1.3) | 0.0 (0) |
| St-Bonaventure | 45.95 N, 72.7 W | 173.8 (-78.6) | 0 (0) | 132.4 (36.9) | 74 (41.7) | 0.0 (0) |
| St-Charles | 46.79 N, 70.96 W | 938.6 (-126.1) | 20.1 (-3.4) | 140.8 (98.9) | 48.6 (25.8) | 0.0 (4.7) |
| St-Fabien | 48.31 N, 68.87 W | 35.3 (0) | 0 (0) | 67.6 (-58.2) | 4.8 (47.4) | 0.0 (10.8) |
| St-Henri | 46.71 N, 71.06 W | 35.9 (-7.3) | 0.2 (0) | 123.6 (-74.7) | 25 (7.6) | 0.0 (74.3) |
| St-Ludger-de-Milot SW | 48.87 N, 71.82 W | 778.9 (-257.7) | 18.8 (0) | 31.3 (230.8) | 0 (16.1) | 0.0 (10.8) |
| St-Raphael | 47.79 N, 64.6 W | 87.9 (0) | 0 (0) | 83 (0) | 11.8 (0) | 0.0 (0) |
| Ste-Marguerite Marie | 48.83 N, 72.18 W | 3562.6 (-558) | 20.7 (0) | 276.3 (163.8) | 14.1 (48.9) | 0.0 (345.3) |
| Total |  | 14411.7 (-2211.3) | 306.6 (-16) | 4425.2 (133.6) | 1547 (1258.5) | 0.0 (818.4) |

In 2019, restored sites represented 22.6 % of areas where peat extraction has ended. Vegetation cover has changed markedly over the years since extracted peat sites were left unrestored. We restricted the comparison to the first 17 years since no restored site was older than this at the time of the study. Based on 146 natural, 221 unrestored and 52 restored sampling sites, *Sphagnum*/moss cover area increased steadily, reaching natural levels ca. 10 y after restoration but not in unrestored sites (Fig. 2; paired samples *t*-test, *t* = 4.5, *p* = 0.0009). Herbaceous strata rapidly reached high percent cover in restored sites, but remained in the natural range in unrestored sites (*t* = 4.7, *p* = 0.0006). Percent cover of other strata was independent of restoration (*p* > 0.05) but Ericaceous shrubs approached natural levels after 17 y.

**Figure 2:**
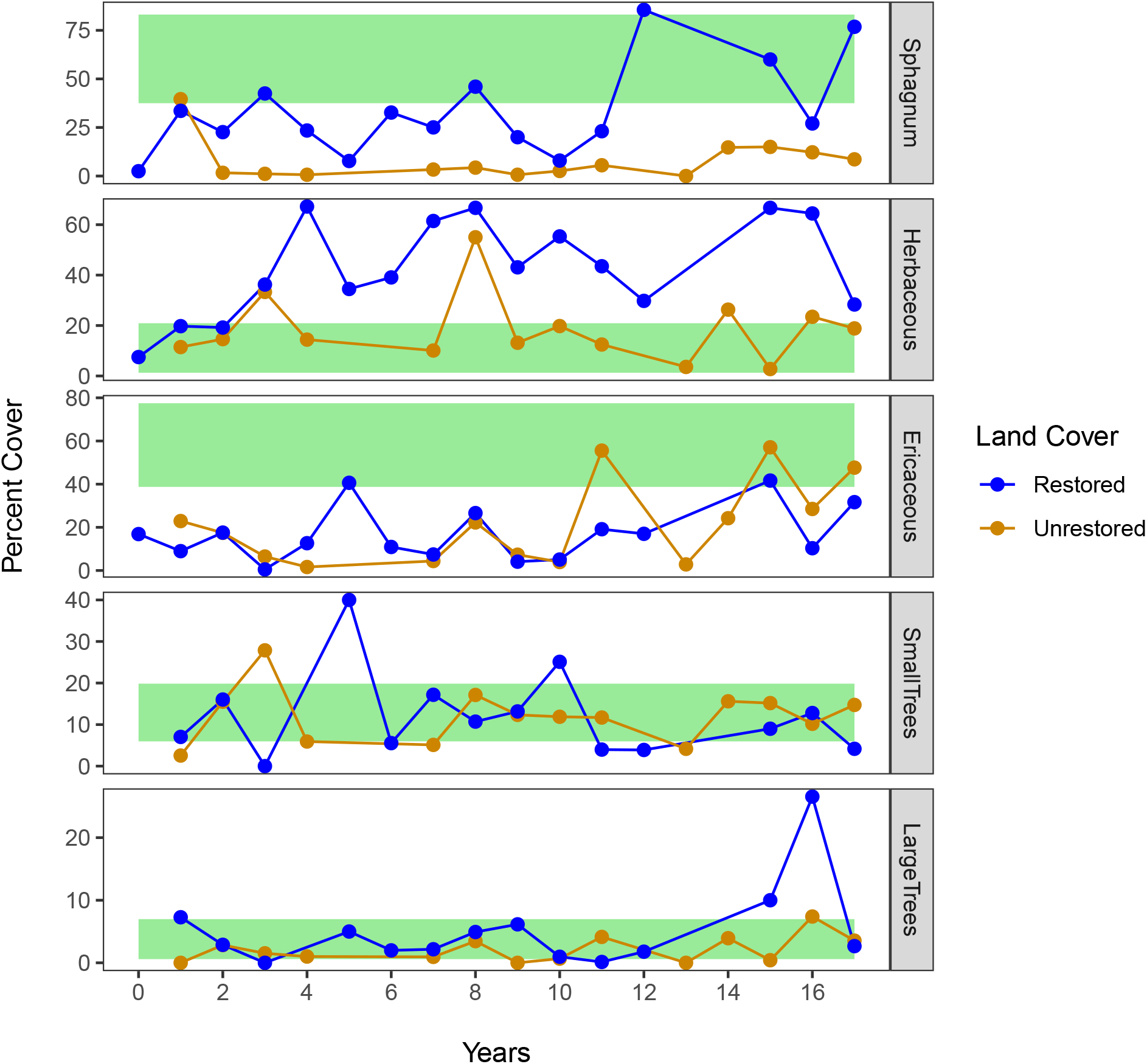
Chronosequence of plant cover across sampled sites. We restricted the comparison to the first 17 years since no restored site was older than this at the time of the study. Green bands represent variability in natural sites (lower and upper quartiles). Peat in sites less than 40 y old was mostly extracted with the vacuum method; older extraction was generally manual (block cut). Only plots with > 1 ha unrestored or restored were retained for this analysis.

### Birds with small home ranges

We recorded 94 bird species, most of which were uncommon (Appendix 1). The 10 most commonly recorded birds were White-throated Sparrow, Common Yellowthroat, Lincoln’s Sparrow, Hermit Thrush, Savannah Sparrow, Palm Warbler, American Crow, American Goldfinch, American Robin, and Alder Flycatcher (by decreasing order), and accounted for 69 % of all individuals recorded. Of those species, only crows have large home ranges. Maximum cumulative numbers of species were very similar among bog types (Fig. 3). However, they were slightly lower in unrestored sites than in natural or restored sites (permutation test, *p* < 0.001).

**Figure 3:**
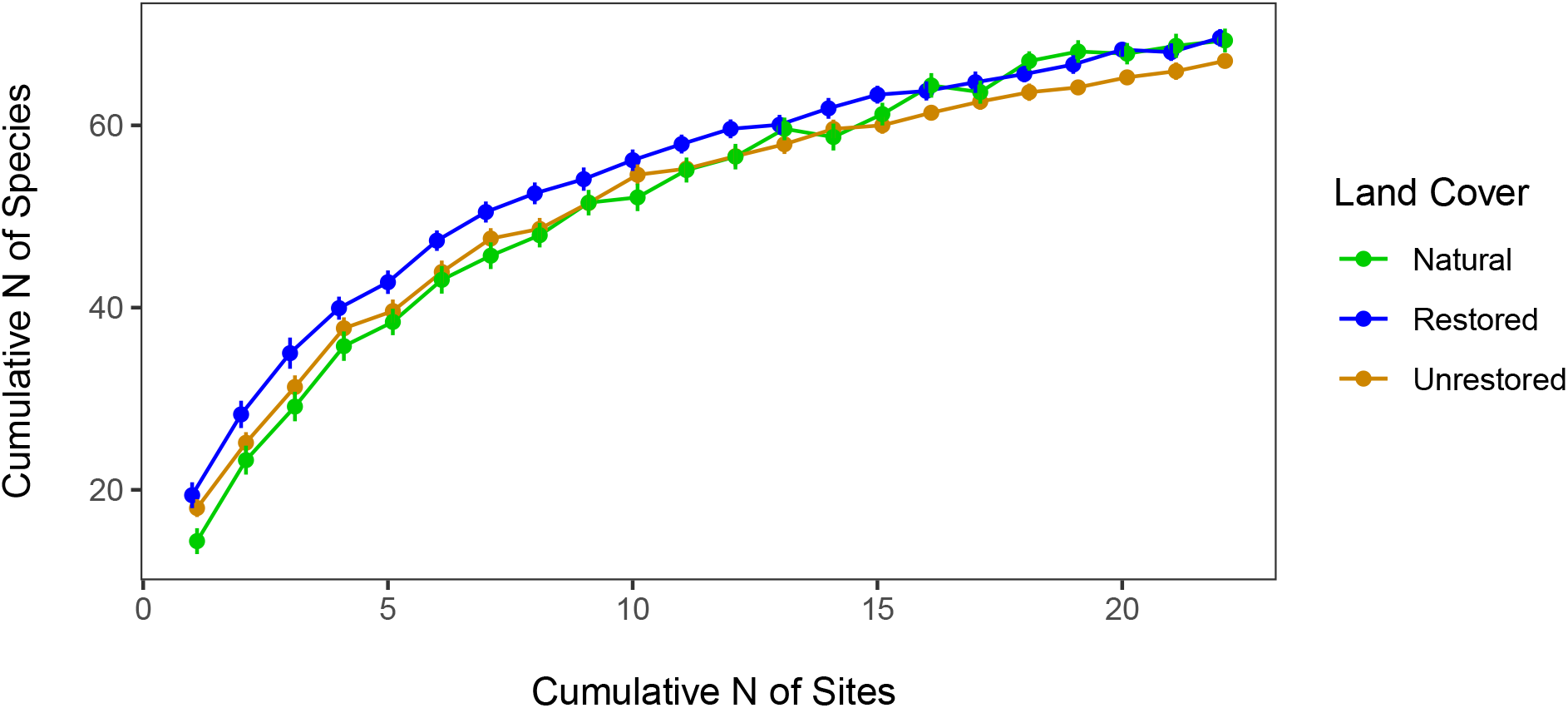
Bird species accumulation curves in natural, unrestored and restored sites. Vertical bars represent bootstrapped standard errors of estimates.

The probability of detection of species with small home ranges during a point count averaged 0.54. However, probabilities of detection varied greatly among species, lowest for American Robin (0.33) and highest for Common Yellowthroat (0.8). Among the 15 species selected for analysis, 6 were significantly associated to one or another of the unrestored, restored, or natural site categories. They were: Alder Flycatcher, American Goldfinch, Palm Warbler, Savannah Sparrow, Song Sparrow, and Yellow-rumped Warbler (Table 3). The habitat associations of 4 of these species appears to have changed with time, as evidenced by their best occupancy model, which included a ‘year × bog status’ interaction. Occupancy of restored sites by Palm and Yellow-rumped Warblers increased over the years, reaching levels similar to those in natural sites (Fig. 4). Occupancy of restored sites by Song and Savannah sparrows also increased, but diverged from their occupancy of natural sites over the course of the study (Fig. 4). We must emphasize that the weight of evidence for a changing association to restored, unrestored or natural sites was overwhelming only in the case of Savannah Sparrow. In the other species shown in Table 3, whether differences in occupancy are constant or changing will only be ascertained by further research.

**Table 3:**
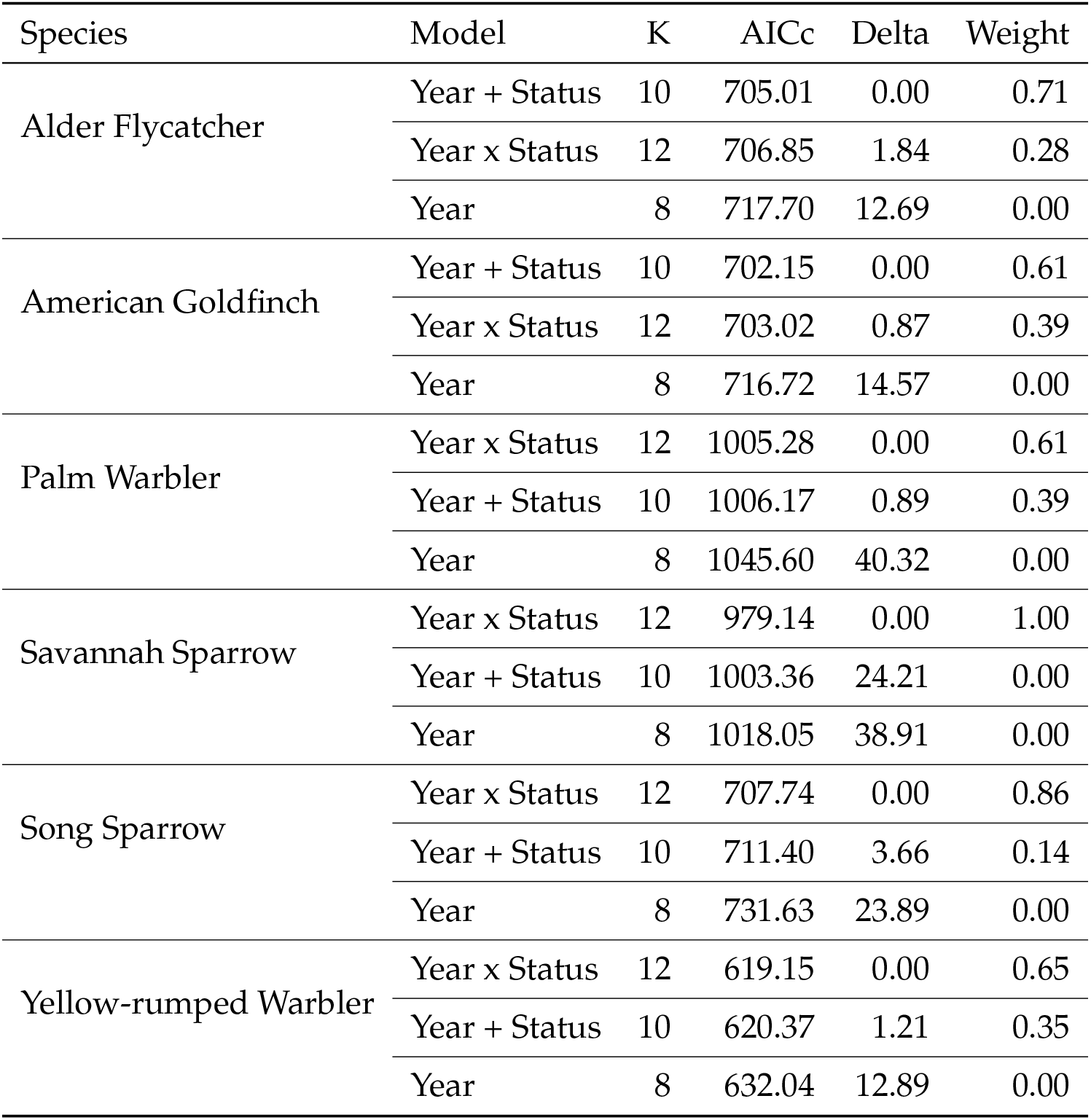
Performance of occupancy models for bird species with small home ranges encountered at least 50 times in the course of the study. Only species for which the null model (year effect only) had a weight less than 5 % (corrected for multiple testing) are shown. K represents the numbers of parameters of each model.

**Figure 4:**
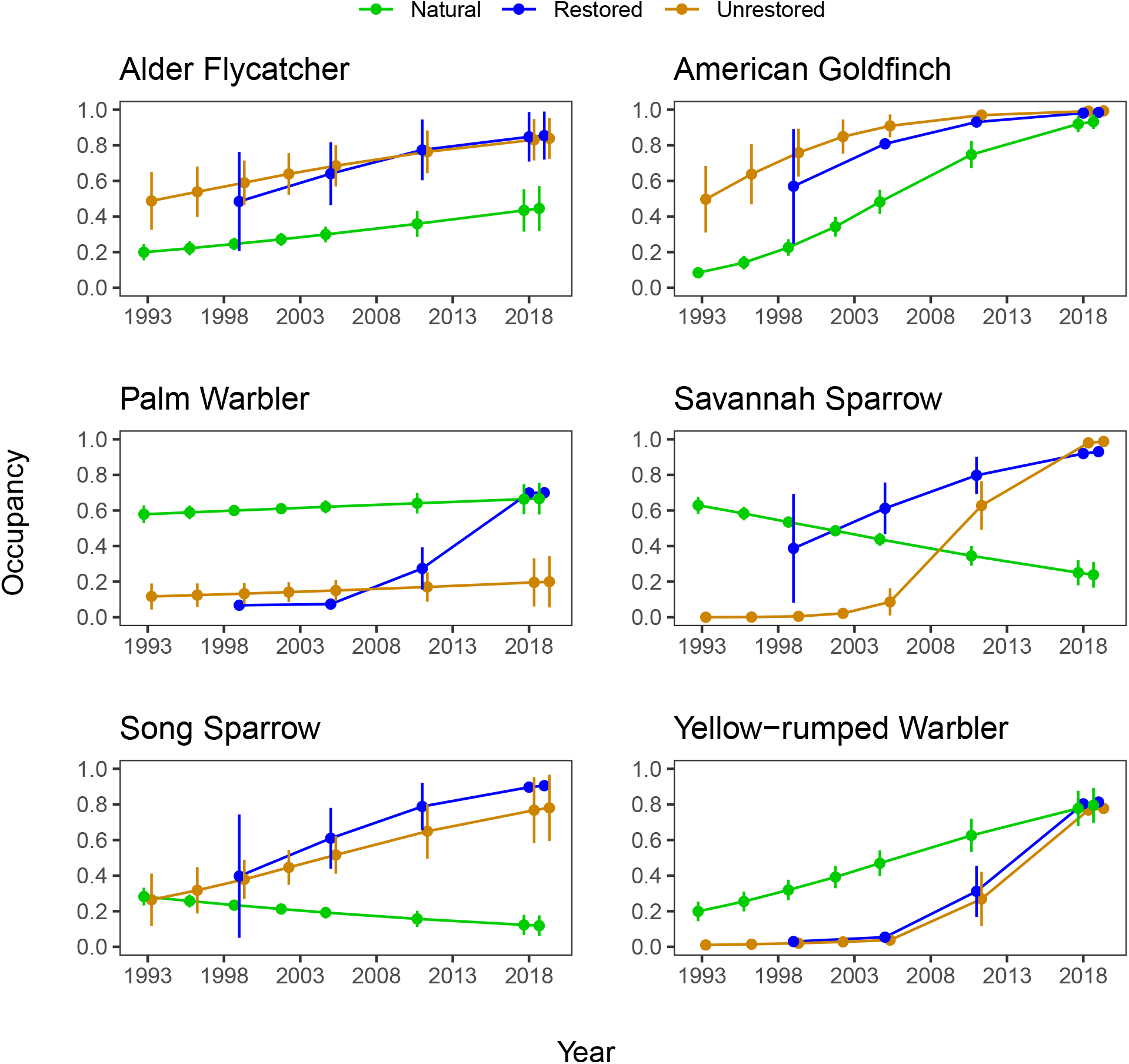
Changing associations to natural, unrestored and restored sites, as determined by occupancy models. Vertical bars represent standard errors of occupancy estimates.

When occupancy of unrestored is compared to restored site occupancy, only the Savannah Sparrow had different occupancies between those sites, as evidenced by the occupancy estimate (logit scale) for bog status effect that did not overlap zero 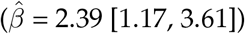.

### Birds with large home ranges

We recorded the following species with large home ranges, or audible from a large distance: American Bittern, American Black Duck, American Crow, American Kestrel, Bank Swallow, Barn Swallow, Belted Kingfisher, Blue Jay, Broad-winged Hawk, Brown-headed Cowbird, Brown Thrasher, Cedar Waxwing, Common Grackle, Common Raven, Eastern Meadowlark, Merlin, Mourning Dove, Northern Harrier, Red-tailed Hawk, Sandhill Crane, Sharp-shinned Hawk, Upland Sandpiper, Tree Swallow, Turkey Vulture, and migrant shorebirds. Migrant shorebirds were composed of 8 species: Greater Yellowlegs, Least Sandpiper, Lesser Yellowlegs, Pectoral Sandpiper, Sanderling, Semipalmated Plover, Semipalmated Sandpiper, and Solitary Sandpiper. Probabilities of detection among species with large home ranges were lowest for migrant shorebirds (0.03) and highest for American Crow (0.45). None of the species responded significantly to the log-transformed areas of ponds, natural, peat-extracted, unrestored and restored sites, after adjusting the type I error risk for multiple testing.

### Bird species assemblages

Species assemblages differed significantly among peat extracted, unrestored and restored sites (Mantel test, *r* = 0.086, *p* < 0.001). After excluding natural sites, species assemblages remained significantly different between restored and unrestored sites (Mantel test, *r* = 0.086, *p* < 0.0006). Species assemblages of unrestored and restored sites were very dissimilar to those of natural sites soon after extraction, but became progressively more similar with increasing years post extraction (Fig. 5; 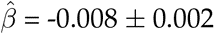, *p* < 0.003).

**Figure 5:**
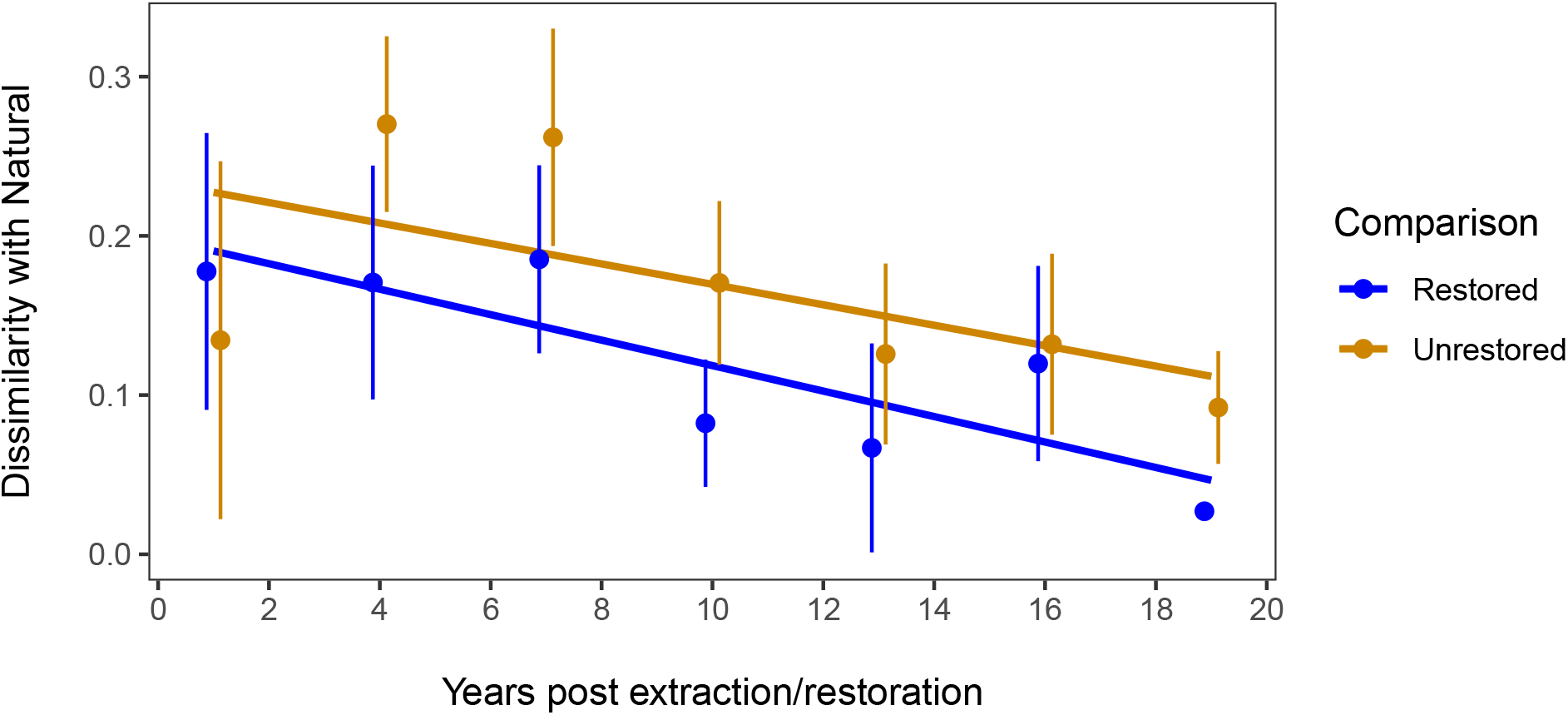
Convergence of bird species assemblages of unrestored and restored sites towards those of natural sites in Eastern canadian bogs, 1993 – 2019.

## Discussion

Bogs subject to peat extraction in Eastern Canada harbour a large number of bird species, but most of them do not occur in those sites regularly, the overall picture resembling a power law (or ‘Pareto’) distribution, as is the case in many phenomena (Newman, 2005) including species abundances (Ulrich et al., 2010). Species diversity was remarkably similar among natural, unrestored, and restored sites, even though unrestored sites yielded slightly fewer species. As in other studies with the present ecosystem (González and Rochefort, 2014; Purre et al., 2020), we show that low vegetation strata can recover in a matter of a few years after peat extraction has ceased, and that this recovery is facilitated by restoration in the case of moss and herbaceous (mostly sedges) strata (Rochefort et al., 2013). Moreover, we added support to the finding by Desrochers et al. (1998) that tree cover overshot in old unrestored sites that of nearby undisturbed sites, a phenomenon linked to the regional establishment of grey birch (*Betula populifolia*) in disturbed bogs (Lavoie and Saint-Louis, 1999). In this context, it is not surprising that site occupancy by certain bird species has changed, since birds tend to respond strongly to vegetation structure, at least at coarse spatial scales (e.g. (Rotenberry, 1985)). But the convergence of bird species assemblages of unrestored and restored sites towards those of natural sites is a less trivial, and encouraging, outcome of this study. If species dissimilarity indices are to be trusted, bog restoration efforts may be on the cusp of yielding bird species assemblages statistically indistinguishable from those of natural sites.

In his classic monograph on boreal birds of Canada, Erskine (1977) provided an in-depth picture of bird assemblages in Canadian boreal bogs, sometimes termed ‘muskeg’. In the 1990s, our research group presented a first outlook on the impact of peat extraction on eastern Canadian birds, with a focus on recolonization (Desrochers et al., 1998). In the natural bogs, we found approximately the same bird species as the ones Erskine found in natural bogs. One notable exception was the Swamp Sparrow, that Erskine depicted as typical of bogs, despite it being rare in the bogs of the present study, and probably more associated to fens. Our 1998 study provided little ground for hope in rebuilding bird assemblages similar to undisturbed bogs. At the time of that study, unrestored vacuum-extracted sites exhibited *Sphagnum* and ericaceous cover well-below levels observed in nearby natural sites, and strikingly different bird species (Desrochers et al., 1998).

Our 1998 study was conducted at a time when bog restoration was still in its infancy, and the ca. 25 years that have elapsed since allow us to provide a more robust assessment of bog recolonization by birds, by documenting long-term changes in bird responses to bog post extraction and restoration. Furthermore, our statistical analyses of single species now benefit from key advances offered by hierarchical modeling that accounts for imperfect detection (Burnham and Anderson, 2002; MacKenzie et al., 2002), thereby providing true occupancy estimates as opposed to occurrence indices such as those used in our earlier work. Only a minority of the species examined exhibited strong differences in occupancy of unrestored, restored and natural sites. Among them, the Savannah Sparrow and the Palm Warbler merit special attention. Savannah Sparrows are found in a variety of open habitats (meadows, crop fields, sand dunes, etc.) and is therefore not a flagship bog species. But the creation of barren sites in bogs, restored or not, seem to have been beneficial for the species, to the point of leading to a greater occupancy of those sites than natural sites which we assumed were optimal for the species, 25 years ago. By contrast, the Palm Warbler, the North American bird most closely associated to bogs in temperate areas (Calmé et al., 2002; Wilson, 2020), seems to have responded well to bog restoration in recent years, while maintaining its occupancy of natural sites.

How does our work compare to other studies of recolonization in restored ecosystems? In the last 20 years, the restoration ecology literature has exploded, but only a small portion of it specifically deals with faunal recolonization (Cross et al., 2019). In most cases, substantial differences have been maintained or even increased between bird assemblages of restored and reference sites. for example, in rehabilitated bauxite mines of northern Australia, Brady and Noske (2010) showed, as in the current study, that species diversity was similar in restored and reference sites, but lasting differences were found in bird species assemblages. In Wisconsin, USA, Hapner et al. (2011) found that restored wetlands were progressively more occupied by oldfield species, and less by wetland-dependent species, likely because of a reduction on the amount of open water (Hapner et al., 2011). Wilson and Bayne (2019) document yet another outcome in Alberta, Canada, where songbird species of reclaimed oil and gas wellsites became more similar to those of nearby mature forest, within 49 years post extraction. The convergence of restored and natural bird species assemblages in our study, and the return of a flagship species, thus falls within the range of documented restoration outcomes, especially if we broaden the definition of ecological restoration to include extensive forestry, where wildlife typically recolonizes clearcut sites after decades of forest succession, in temperate as well as tropical forests (Acevedo-Charry and Aide, 2019, and references therein).

The restoration of raised bogs has been underway not only in North America, but also in Europe, most notably in Baltic countries (Karofeld et al., 2017). From the ornithological point of view, advances have been made to restore bogs for a few selected species such as the Golden Plover (*Pluvialis apricaria*), whose regional decline has been attributed in part to the loss of nesting habitat (Minayeva et al., 2017). Finland presents a particular case, with over 300 km^2^ of forested peatlands restored by blocking drainage ditches following agriculture and forestry (Alsila et al., 2021). To our knowledge, the only quantitative assessment of the effectiveness of peatland restoration for birds in northern Europe comes from Alsila et al. (2021), who found no response to restoration by peatland specialist birds, based on annual surveys just before restoration and up to 8 y post restoration. One key difference in bird life between European and North American bogs, at least in regions where peat extraction occurs, is the prevalence of songbirds and the paucity of ducks and shorebirds in the latter (Desrochers and van Duinen, 2006). It is therefore likely that bog restoration success in northern Europe will hinge on the ability to rebuild ponds in sites where peat has been extracted.

### Moving goal posts

Recolonization by flagship bird species may mark the success of ecological restoration from an ornithological perspective. But more generally, what marks the success of ecological restoration? The question has been debated for some time (Ruiz-Jaen and Mitchell Aide, 2005) and answers will undoubtedly vary because of the diversity of stated objectives. Even in the relatively simple case of vegetation recovery, proponents of ecological restoration are daunted by the challenge of defining goals in a constantly changing environment, with moving goal posts. What amount of ecological variation in time and space can be considered desirable? To compound the problem, the “goal posts” of site recolonization by mobile animals such as birds can be moving independently of abiotic factors, vegetation structure, policy targets, etc., because of regional and sometimes faraway factors, in the case of long-distance migrants. This problem is well illustrated with Savannah Sparrows, whose “reference” level of abundance has been declining in the last 25 years, for reasons probably unrelated to the fortunes of bogs under production. Conversely, the “reference” level of abundance of Sandhill Cranes has moved to vanishingly small to very high in the province of Quebec since the onset of the present study, again with no apparent link to peat extraction. So one may ask, how many Savannah Sparrows or Sandhill Cranes would be considered ideal in restored bogs?

Besides the “moving goal posts” problem, ecological restoration may suffer from an overly restrictive focus on reference ecosystems, without an openness to unexpected opportunity. In the Czech Republic, Šálek (2012) discovered that restoration efforts actually *reduced* the value of post mining sites for birds, because they focused on the establishment of trees, to the detriment of barren grounds which were more beneficial for regionally-endangered birds. In a similar vein, vacuum-harvesting of peat, with no subsequent restoration, has inadvertently created novel habitats for species of concern in the region concerned by the present study, such as Savannah Sparrow (documented above), Vesper Sparrow, Killdeer and Common Nighthawk. Populations of those species have declined precipitously in Canada since 1970 (by 64 %, 61 %, and 49 % respectively; (NABCI, 2019)). Those declining species are associated to barren sites, not to dense ericaceous shrubs. We observed the latter three species frequently in the extracted bogs of this study, but they were seldom, if at all, observed in undisturbed bogs (Calmé and Desrochers, 2000, 1999). Thus for certain species, the lack of intervention may lead to more desirable outcomes than active restoration.

If land cover management and restoration success result in the conservation of sizable areas of undisturbed and restored peatlands, those habitats may be able to persist longer than surrounding upland habitat, because of their hydrological resilience (Waddington et al. 2015). Restored and natural bogs will continue to provide opportunity to measure restoration, recolonization, and perhaps unexpected opportunity.

## Acknowledgments

This research was funded by Natural Science and Engineering Research Council of Canada’s, the Canadian Sphagnum Peat Moss Association and its numerous corporate members, as well as a Wildlife Habitat Canada grant to André Desrochers. Thanks to Alexandre Rivard, Bertrand LeGrand, Bruno Drolet, Christian Marcotte, Christine Renaud, Fanny Senez-Gagnon, Jean-François Ouellet, Jean-François Rousseau, Jean-Pierre Savard, Nathalie Pelletier, and Stéphanie Haddad for their participation in bird point counts. Stéphanie Boudreau provided helpful criticism on an earlier version of this article.

## Appendix

**Table 4:** List of species, by decreasing number of records.

| English name | Scientific name | Natural | Unrestored | Restored | Total |
| --- | --- | --- | --- | --- | --- |
| Common Yellowthroat | *Geothlypis trichas* | 1895 | 228 | 161 | 2284 |
| White-throated Sparrow | *Zonotrichia albicollis* | 1599 | 299 | 188 | 2086 |
| Lincoln's Sparrow | *Melospiza lincolnii* | 1260 | 103 | 95 | 1458 |
| Savannah Sparrow | *Passerculus sandwichensis* | 879 | 48 | 131 | 1058 |
| Palm Warbler | *Setophaga palmarum* | 966 | 20 | 38 | 1024 |
| Hermit Thrush | *Catharus guttatus* | 710 | 121 | 129 | 960 |
| American Crow | *Corvus brachyrhynchos* | 423 | 87 | 89 | 599 |
| Alder Flycatcher | *Empidonax alnorum* | 261 | 101 | 115 | 477 |
| American Goldfinch | *Spinus tristis* | 225 | 130 | 94 | 449 |
| Song Sparrow | *Melospiza melodia* | 244 | 96 | 90 | 430 |
| American Robin | *Turdus migratorius* | 246 | 88 | 82 | 416 |
| Magnolia Warbler | *Setophaga magnolia* | 260 | 45 | 35 | 340 |
| Ruby-crowned Kinglet | *Regulus calendula* | 245 | 26 | 29 | 300 |
| Red-winged Blackbird | *Agelaius phoeniceus* | 179 | 44 | 67 | 290 |
| Yellow-rumped Warbler | *Setophaga coronata* | 231 | 16 | 24 | 271 |
| Red-eyed Vireo | *Vireo olivaceus* | 96 | 47 | 50 | 193 |
| Purple Finch | *Haemorhous purpureus* | 128 | 11 | 12 | 151 |
| Cedar Waxwing | *Bombycilla cedrorum* | 101 | 20 | 10 | 131 |
| Swainson's Thrush | *Catharus ustulatus* | 82 | 19 | 20 | 121 |
| Veery | *Catharus fuscescens* | 49 | 39 | 27 | 115 |
| Common Grackle | *Quiscalus quiscula* | 86 | 10 | 16 | 112 |
| American Redstart | *Setophaga ruticilla* | 56 | 25 | 28 | 109 |
| Black-capped Chickadee | *Poecile atricapillus* | 63 | 19 | 20 | 102 |
| Swamp Sparrow | *Melospiza georgiana* | 8 | 31 | 56 | 95 |
| Blue Jay | *Cyanocitta cristata* | 65 | 4 | 14 | 83 |
| Wilson's Snipe | *Gallinago delicata* | 31 | 18 | 33 | 82 |
| Nashville Warbler | *Leiothlypis ruficapilla* | 45 | 12 | 23 | 80 |
| Common Raven | *Corvus corax* | 44 | 10 | 20 | 74 |
| Killdeer | *Charadrius vociferus* | 31 | 27 | 11 | 69 |
| American Black Duck | *Anas rubripes* | 54 | 6 | 5 | 65 |
| Northern Harrier | *Circus hudsonius* | 47 | 10 | 7 | 64 |
| Tree Swallow | *Tachycineta bicolor* | 52 | 4 | 7 | 63 |
| Eastern Kingbird | *Tyrannus tyrannus* | 54 | 3 | 5 | 62 |
| Vesper Sparrow | *Poocetes gramineus* | 11 | 22 | 26 | 59 |
| Philadelphia Vireo | *Vireo philadelphicus* | 32 | 12 | 13 | 57 |
| Northern Flicker | *Colaptes auratus* | 19 | 11 | 21 | 51 |
| Winter Wren | *Troglodytes hiemalis* | 37 | 7 | 6 | 50 |
| Chestnut-sided Warbler | *Setophaga pensylvanica* | 22 | 13 | 9 | 44 |
| Brown-headed Cowbird | *Molothrus ater* | 30 | 7 | 6 | 43 |
| Dark-eyed Junco | *Junco hyemalis* | 30 | 5 | 5 | 40 |
| Ovenbird | *Seiurus aurocapilla* | 28 | 10 | 2 | 40 |
| Golden-crowned Kinglet | *Regulus satrapa* | 36 | 2 | 1 | 39 |
| Mourning Dove | *Zenaida macroura* | 31 | 6 | 0 | 37 |
| Pine Siskin | *Spinus pinus* | 27 | 3 | 7 | 37 |
| Boreal Chickadee | *Poecile hudsonicus* | 31 | 1 | 2 | 34 |
| Upland Sandpiper | *Bartramia longicauda* | 34 | 0 | 0 | 34 |
| Mallard | *Anas platyrhynchos* | 24 | 2 | 7 | 33 |

Table 4: List of species, by decreasing number of records. (continued)
| English name | Scientific name | Natural | Unrestored | Restored | Total |
| --- | --- | --- | --- | --- | --- |
| Red-breasted Nuthatch | *Sitta canadensis* | 20 | 4 | 6 | 30 |
| Yellow-bellied Flycatcher | *Empidonax flaviventris* | 23 | 3 | 3 | 29 |
| Black-and-white Warbler | *Mniotilta varia* | 22 | 3 | 0 | 25 |
| Canada Goose | *Branta canadensis* | 17 | 1 | 7 | 25 |
| Common Loon | *Gavia immer* | 16 | 6 | 3 | 25 |
| White-winged Crossbill | *Loxia leucoptera* | 22 | 2 | 1 | 25 |
| American Woodcock | *Scolopax minor* | 12 | 9 | 2 | 23 |
| Least Flycatcher | *Empidonax minimus* | 3 | 10 | 8 | 21 |
| Tennessee Warbler | *Leiothlypis peregrina* | 7 | 1 | 13 | 21 |
| Olive-sided Flycatcher | *Contopus cooperi* | 18 | 1 | 1 | 20 |
| Blue-headed Vireo | *Vireo solitarius* | 12 | 6 | 1 | 19 |
| Yellow Warbler | *Setophaga petechia* | 4 | 10 | 5 | 19 |
| Bobolink | *Dolichonyx oryzivorus* | 9 | 3 | 6 | 18 |
| Black-throated Green Warbler | *Setophaga virens* | 11 | 4 | 1 | 16 |
| Barn Swallow | *Hirundo rustica* | 8 | 2 | 5 | 15 |
| Evening Grosbeak | *Coccothraustes vespertinus* | 10 | 5 | 0 | 15 |
| Gray Catbird | *Dumetella carolinensis* | 12 | 0 | 3 | 15 |
| American Bittern | *Botaurus lentiginosus* | 4 | 0 | 10 | 14 |
| Bank Swallow | *Riparia riparia* | 7 | 0 | 5 | 12 |
| Short-eared Owl | *Asio flammeus* | 12 | 0 | 0 | 12 |
| No birds | *Non avium* | 5 | 0 | 5 | 10 |
| Sandhill Crane | *Antigone canadensis* | 6 | 2 | 2 | 10 |
| Brown Thrasher | *Toxostoma rufum* | 0 | 0 | 9 | 9 |
| Common Nighthawk | *Chordeiles minor* | 5 | 1 | 3 | 9 |
| Great Blue Heron | *Ardea herodias* | 6 | 2 | 1 | 9 |
| Mourning Warbler | *Geothlypis philadelphia* | 5 | 3 | 1 | 9 |
| Rusty Blackbird | *Euphagus carolinus* | 8 | 1 | 0 | 9 |
| Green-winged Teal | *Anas crecca* | 6 | 0 | 2 | 8 |
| Herring Gull | *Larus argentatus* | 5 | 0 | 3 | 8 |
| Northern Waterthrush | *Parkesia noveboracensis* | 6 | 1 | 0 | 7 |
| Ruby-throated Hummingbird | *Archilochus colubris* | 5 | 2 | 0 | 7 |
| Wilson's Warbler | *Cardellina pusilla* | 2 | 1 | 4 | 7 |
| Blackpoll Warbler | *Setophaga striata* | 5 | 1 | 0 | 6 |
| Chipping Sparrow | *Spizella passerina* | 2 | 2 | 2 | 6 |
| Ring-billed Gull | *Larus delawarensis* | 4 | 1 | 0 | 5 |
| Yellow-bellied Sapsucker | *Sphyrapicus varius* | 5 | 0 | 0 | 5 |
| American Pipit | *Anthus rubescens* | 3 | 1 | 0 | 4 |
| Bay-breasted Warbler | *Setophaga castanea* | 4 | 0 | 0 | 4 |
| Black-billed Cuckoo | *Coccyzus erythrophthalmus* | 2 | 0 | 2 | 4 |
| Cape May Warbler | *Setophaga tigrina* | 1 | 2 | 1 | 4 |
| Cliff Swallow | *Petrochelidon pyrrhonota* | 4 | 0 | 0 | 4 |
| Fox Sparrow | *Passerella iliaca* | 4 | 0 | 0 | 4 |
| Osprey | *Pandion haliaetus* | 3 | 0 | 1 | 4 |
| Pied-billed Grebe | *Podilymbus podiceps* | 0 | 0 | 4 | 4 |
| Pileated Woodpecker | *Dryocopus pileatus* | 2 | 0 | 2 | 4 |
| American Wigeon | *Mareca americana* | 0 | 0 | 3 | 3 |
| Canada Warbler | *Cardellina canadensis* | 2 | 1 | 0 | 3 |
| Double-crested Cormorant | *Phalacrocorax auritus* | 1 | 2 | 0 | 3 |
| Great Crested Flycatcher | *Myiarchus crinitus* | 3 | 0 | 0 | 3 |
| Least Sandpiper | *Calidris minutilla* | 2 | 0 | 1 | 3 |

Table 4: List of species, by decreasing number of records. (continued)
| English name | Scientific name | Natural | Unrestored | Restored | Total |
| --- | --- | --- | --- | --- | --- |
| Northern Parula | *Setophaga americana* | 1 | 2 | 0 | 3 |
| Scarlet Tanager | *Piranga olivacea* | 2 | 1 | 0 | 3 |
| Sharp-shinned Hawk | *Accipiter striatus* | 2 | 0 | 1 | 3 |
| Spotted Sandpiper | *Actitis macularius* | 2 | 0 | 1 | 3 |
| American Kestrel | *Falco sparverius* | 2 | 0 | 0 | 2 |
| Bald Eagle | *Haliaeetus leucocephalus* | 1 | 1 | 0 | 2 |
| Belted Kingfisher | *Megasceryle alcyon* | 2 | 0 | 0 | 2 |
| Canada Jay | *Perisoreus canadensis* | 2 | 0 | 0 | 2 |
| Northern Hawk Owl | *Surnia ulula* | 2 | 0 | 0 | 2 |
| Red-tailed Hawk | *Buteo jamaicensis* | 2 | 0 | 0 | 2 |
| Rose-breasted Grosbeak | *Pheucticus ludovicianus* | 2 | 0 | 0 | 2 |
| Ruffed Grouse | *Bonasa umbellus* | 2 | 0 | 0 | 2 |
| Solitary Sandpiper | *Tringa solitaria* | 0 | 1 | 1 | 2 |
| Black-throated Blue Warbler | *Setophaga caerulescens* | 1 | 0 | 0 | 1 |
| Blackburnian Warbler | *Setophaga fusca* | 1 | 0 | 0 | 1 |
| Clay-colored Sparrow | *Spizella pallida* | 1 | 0 | 0 | 1 |
| Common Tern | *Sterna hirundo* | 0 | 0 | 1 | 1 |
| Downy Woodpecker | *Dryobates pubescens* | 0 | 0 | 1 | 1 |
| Eastern Bluebird | *Sialia sialis* | 1 | 0 | 0 | 1 |
| Eastern Meadowlark | *Sturnella magna* | 0 | 1 | 0 | 1 |
| Eastern Wood-Pewee | *Contopus virens* | 1 | 0 | 0 | 1 |
| Great Black-backed Gull | *Larus marinus* | 0 | 1 | 0 | 1 |
| Great Gray Owl | *Strix nebulosa* | 1 | 0 | 0 | 1 |
| Greater Yellowlegs | *Tringa melanoleuca* | 0 | 0 | 1 | 1 |
| Hairy Woodpecker | *Dryobates villosus* | 0 | 0 | 1 | 1 |
| Horned Lark | *Eremophila alpestris* | 1 | 0 | 0 | 1 |
| Lesser Yellowlegs | *Tringa flavipes* | 0 | 0 | 1 | 1 |
| Merlin | *Falco columbarius* | 0 | 0 | 1 | 1 |
| Northern Mockingbird | *Mimus polyglottos* | 1 | 0 | 0 | 1 |
| Pectoral Sandpiper | *Calidris melanotos* | 1 | 0 | 0 | 1 |
| Ring-necked Duck | *Aythya collaris* | 1 | 0 | 0 | 1 |
| Sanderling | *Calidris alba* | 1 | 0 | 0 | 1 |
| Spruce Grouse | *Falcipennis canadensis* | 1 | 0 | 0 | 1 |
| Turkey Vulture | *Cathartes aura* | 1 | 0 | 0 | 1 |
| White-breasted Nuthatch | *Sitta carolinensis* | 1 | 0 | 0 | 1 |
| White-crowned Sparrow | *Zonotrichia leucophrys* | 1 | 0 | 0 | 1 |

